# Pair consensus decoding improves accuracy of neural network basecallers for nanopore sequencing

**DOI:** 10.1101/2020.02.25.956771

**Authors:** Jordi Silvestre-Ryan, Ian Holmes

## Abstract

Nanopore technology allows for direct sequencing of individual DNA duplexes. However, its higher error rate compared to other sequencing methods has limited its application in situations where deep coverage is unavailable, such as detection of rare variants or characterization of highly polymorphic samples. In principle, 2X coverage is available even for single duplexes, using Oxford Nanopore Technologies’ 1D^2^ protocol or related methods which sequence both strands of the duplex consecutively. Using both strands should improve accuracy; however, most neural network basecaller architectures are designed to operate on single strands. We have developed a general approach for improving accuracy of 1D^2^ and related protocols by finding the consensus of two neural network basecallers, by combining a constrained profile-profile alignment with a heuristic variant of beam search. When run on a basecalling neural network we trained, our consensus algorithm improves median basecall accuracy from 86.2% (for single-read decoding) to 92.1% (for pair decoding). Our software can readily be adapted to work with the output of other basecallers, such as the recently released Bonito basecaller. Although Bonito operates only on individual strands and was not designed to leverage the 1D^2^ protocol, our method lifts its median accuracy from 93.3% to 97.7%, more than halving the median error rate. This surpasses the maximum accuracy achievable with Guppy, an alternate basecaller which was designed to include pair decoding of 1D^2^ reads. Our software PoreOver, including both our neural network basecaller and our consensus pair decoder (which can be separably applied to improve other basecallers), is implemented in Python 3 and C++11 and is freely available at https://github.com/jordisr/poreover.

## Introduction

While the use of protein nanopores to sequence DNA has been explored for many years, the technology was not commercialized until 2014 when Oxford Nanopore Technologies (ONT) released their MinION device[1]. Nanopore sequencing can generate long reads in the tens or hundreds of kilobases—including direct readout of modified bases—and can be run on a highly portable platform, enabling sequencing in locations as remote as the International Space Station[2]. However, the relatively high error rate of nanopore sequencing has limited some of its applications and spurred the development of new chemistries and basecalling methods [3].

Nanopore sequencing works by threading a single strand of DNA through a protein nanopore embedded in a synthetic membrane. The DNA bases block the pore, perturbing the ionic current flowing through the pore. Machine learning methods are needed to learn this complex, sequence-dependent nanopore current function and deduce the sequence of nucleotides that passed through the pore. To do this, basecalling has made heavy use of probabilistic models: the early ONT basecaller used hidden Markov models, as did the community basecaller Nanocall [4]. Mirroring a shift in other areas of bioinformatics, a deep learning approach was adopted for higher accuracy, both by ONT and community basecallers such as DeepNano[5] and Basecrawller[6].

As an added complication, each nucleotide spends a random length of time in the pore, and so may generate zero, one, or many current samples during its transition. These early recurrent neural network basecallers used a segmentation step in preprocessing to cluster measurements into discrete “events”, which were then fed into the neural network. In this aspect, basecalling shares similarities with the task of speech recognition, namely a numeric input that must be segmented and labeled with a discrete number of categories: phonemes or nucleotides. Connectionist Temporal Classification (CTC) [7] was developed as a way to train recurrent neural networks to simultaneously classify and segment such a time series. Using dynamic programming, a loss could be calculated between the deduced and the true sequence labeling; this loss function can then be minimized via gradient descent. The community basecaller Chiron[8] applied CTC to nanopore basecalling, yielding competitive results [9], while ONT incorporated CTC-style models into both production and research basecallers.

The outputs of the neural networks that are trained with CTC loss functions can be interpreted as weighted transitions in a finite state machine, not unlike a profile HMM[10]. Thus, the neural network output defines a probability distribution *P*(*ℓ*|*y*) over possible basecalled sequences *ℓ* given the read *y*. For any given sequence *ℓ*, this probability is computed using the Forward algorithm. By analogy to HMMs, the task of finding the modal sequence of this distribution is termed “decoding”. While perfectly optimal decoding requires an intractable exhaustive search over sequences, in practice heuristic algorithms (such as beam search or Viterbi search) can be used to find reasonably good solutions.

The related problem of “consensus decoding” arises when multiple reads {*y_n_*} are derived from the same underlying sequence *ℓ*. Basecalling then yields multiple profiles. We seek to find the single sequence that maximizes *P*(*ℓ*|{*y_n_*}); under a flat prior *P*(*ℓ*) and the assumption that the reads are independent, this will be the sequence that maximizes the product of Forward probabilities Π_*n*_ *P*(*ℓ*|*y_n_*), motivating the reframing of this problem as an exercise in profile-profile alignment[11]. Pairwise consensus basecalling is particularly relevant to ONT’s 1D^2^ sequencing protocol, in which special DNA adapters are used such that after the template DNA strand passes through the pore, its complementary strand very often follows.

To this end we have developed a beam search decoding algorithm for the pair decoding of two reads, making use of a constrained dynamic programming alignment envelope heuristic to speed calculations by focusing on areas of each read which are likely to represent the same sequence. We test this with our own basecalling software PoreOver, which implements a CTC-style recurrent neural network basecaller and associated CTC decoding algorithms. Using the neural network output from PoreOver, our consensus algorithm yields a median 6% improvement in accuracy on a test set of 1D^2^ data. We adapt our software to work with probabilities generated from the ONT production basecaller Guppy as well as Bonito, a recently released research basecaller, both of which make use of CTC-style models. We apply our consensus decoding to output from Bonito and find it achieves the highest median consensus accuracy of 97.7%, surpassing the 1D^2^ method in Guppy.

We additionally assess how the choice of decoding algorithm affects single read accuracy and find that the use of a beam search only slightly improves basecalling accuracy compared with a best-path Viterbi search on our simplified CTC model. However, when applied to the “flip-flop” CTC model used in the latest generation of Oxford Nanopore basecallers we find that more costly beam search does not improve on the best path Viterbi solution.

## Methods

### Basecalling with connectionist temporal classification

Under CTC, the output of the neural network defines a probability distribution over possible labelings of the input, a “labeling” in this case representing the DNA sequence that passed through the pore. CTC uses a differentiable loss function calculated with dynamic programming to calculate the probability of a given labeling. Using gradient descent, the network is trained to maximize the probability of the correct sequence.

The final softmax layer of the neural network outputs *y*, a 5 × *T* matrix that specifies the probability of emitting each base plus a gap character at each step of the input. Let *y*(*t, c*) be the probability of the character 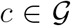 at time *t*, with 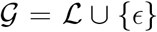, where characters in 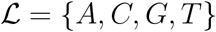 represent bases passing through the pore and *ϵ* is a gap or blank character representing no change in the pore. The neural network output can thus be interpreted as a linear hidden Markov model [11], with the softmax probabilities corresponding to emission probabilities in this HMM. The probability of a given path through this profile 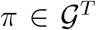 is just the product of the individual probabilities

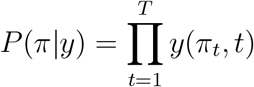

Under this model, sequences longer than *T* have zero probability.

Each gapped path *π* can be mapped to an ungapped label sequence *ℓ* by a function 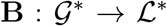, which simply removes the gap characters. This is a simplifed version of the path-to-label mapping used by the canonical CTC model [7]; unlike the original, our version does not merge repeated label characters (so the path `A-CCG--T` would result in the label `ACCGT` rather than `ACGT`). For a given label sequence *ℓ*, we want to find the probability that *ℓ* was emitted by *y*, *P*(*ℓ*|*y*). The probability of a label sequence is the sum of probabilities of all paths consistent with it, essentially marginalizing over all possible positions of gaps:

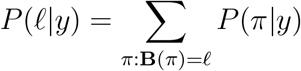

This probability can be efficiently computed by dynamic programming with a Forward algorithm[10]. We define the Forward probability of a label as *α*(*t, s*), the probability that the first *s* characters of *ℓ* having been emitted by position *t* of the underlying HMM. The core recursion is,

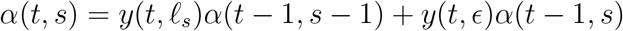

terminated by the base case *α*(0, 0) = 1. From this matrix we can easily read out the probability of the full sequence, *P*(*ℓ*|*y*) = *α*(*T*,|*ℓ*|).

### Decoding the basecaller output

For basecalling we want to find the best label, 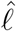,

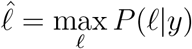

Borrowing HMM terminology, this task is referred to as decoding. While there is not an efficient general algorithm for this optimization, various heuristic search algorithms can be used to find high probability sequences. Here we focus on two approximate methods, finding the (1) Viterbi best path, and (2) a beam search.

While finding the best label is intractable, a simpler approach is to instead find the single best path:

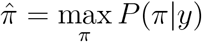

Thus, the Viterbi solution is 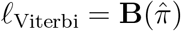. In our CTC model, this is equivalent to taking the argmax of each output in the time series and removing the gaps.

An alternative heuristic search method is beam search, which has been used extensively in the decoding of neural networks, including CTC models[12]. Beam search iterates through the output *y*, keeping a fixed-size list (or ‘beam’) of the best solutions. The size of this list is parameter termed the beam width, and represented with *W*. The algorithm is the following: for each iteration *t* in {1..*T*}, update its probability at time *t* using the forward recursions. Next, extend each label in the beam by each character in the alphabet {*A, C, G, T*} and calculate the corresponding forward probabilities. Finally, prune the beam down to the *W* labels with the highest probabilities. By increasing the beam width *W*, more solutions are tracked at every iteration.

### Neural network architecture and training

Our network consists of a single convolutional layer followed by three bidirectional GRU[13] layers (Figure 1B). Output is passed through a softmax function to yield probabilities for each nucleotide plus a gap character, {*A, C, G, T, ϵ*}. Under this model, the gap character represents no change in the pore, and so the output probability trace consists of peaks of probability as each nucleotide passes through the pore, followed by stretches with high gap probability (Figure 1A).

**Figure 1:**
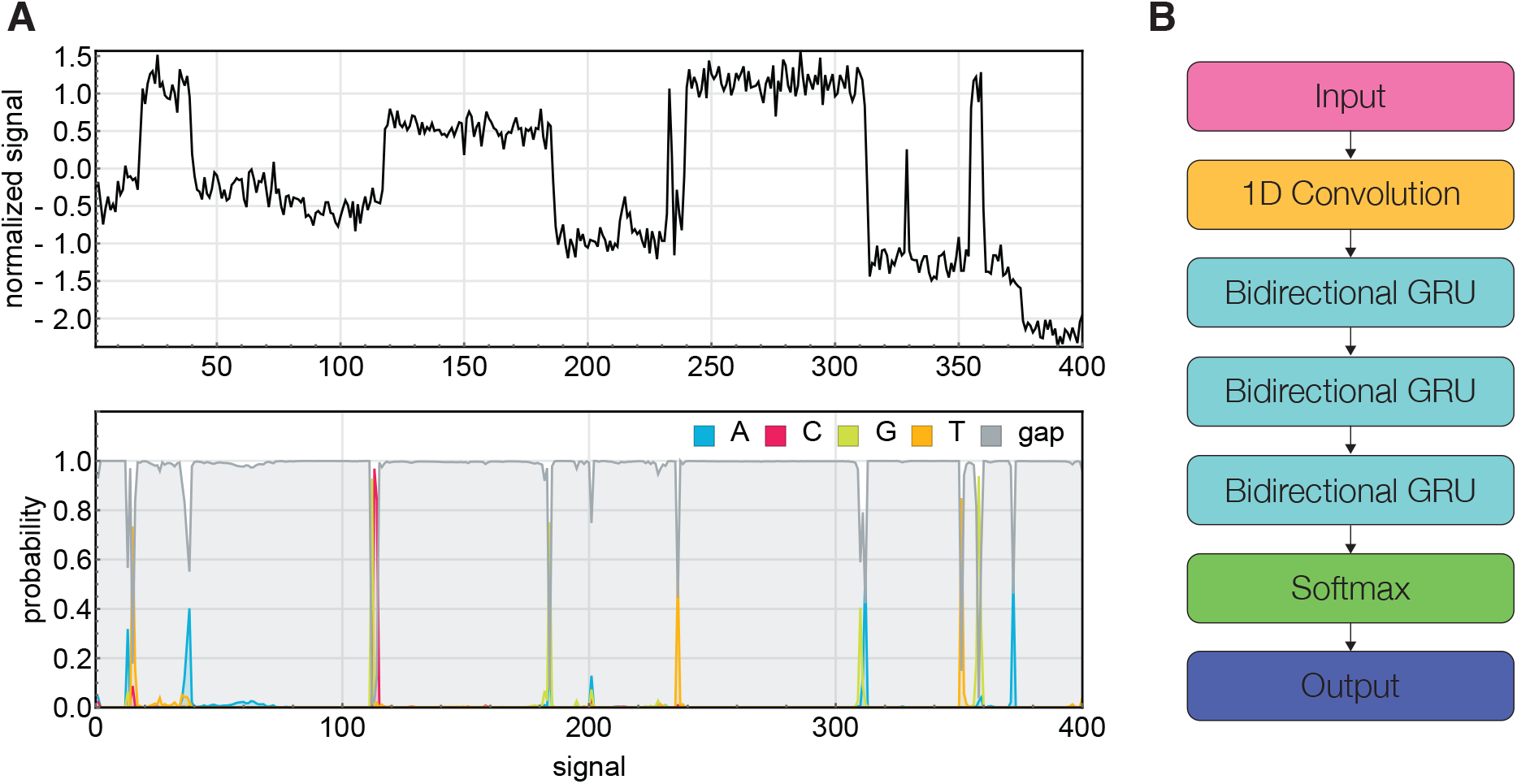
(A) The time series of current signal is translated to probabilities at of each base plus a gap character. The gap represents no change in the pore, so we see a high gap probability punctuated by peaks of probability for each nucleotide. (B) An RNN architecture for basecalling, with a single convolutional layer followed by three stacked bidirectional GRU layers. The softmax output is fed into a CTC loss function, and minimized during training.

Given that training organism influences the generalizability of the network[6], we sought to include a broad taxonomic diversity. A training set of R9.4 reads was assembled from a sample of 10,000 human reads from the nanopore whole genome sequencing consortium[14] and 10,000 microbial reads from the Zymo mock community[15] spanning 8 bacterial and 2 yeast species.

Reads were re-bascalled with the Guppy basecaller and mapped back to reference genomes. Tombo^1^ was used to re-align the raw signal to the reference sequence and correct previous basecalling errors. These corrected reads were split into chunks of 1000 measurements and used as the training set. A fraction of the data was held out as a test set and used to evaluate the performance of our model during training. The neural network was trained for two epochs using the Adam optimizer [16] (learning rate of 0.001) to minimize CTC loss.

## Results

### Consensus decoding of 1D^2^ reads with a banded 2D beam search

Given the probabilistic formulation of decoding CTC models, it is natural to extend decoding for the task of consensus, finding the single sequence that was most likely to be generated by multiple reads. We focus here on the task of decoding a pair of reads, particularly relevant to the 1D^2^ sequencing protocol, which yields separate basecaller probability profiles for both template and complement strands.

Assuming the independence of each read, the best label is given by the consensus probability *P*(*ℓ*|*y*_1_, *y*_2_) = *P*(*ℓ*|*y*_1_)*P*(*ℓ*|*y*_2_)

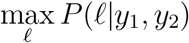

We extend our beam search to work in two dimensions over the *T*_1_ × *T*_2_ space of both reads. At each *t*_1_ ∈ 1..*T*_1_ we update the forward probabilities for labels in the beam at *t*_1_ and for *t*_2_ ∈ 1..*T*_2_. Each label in the beam is assigned a score equal to the forward probability from read 1 at *t*_1_ times the maximum forward probability from read 2 in the range 1..*T*_2_. Finally, the beam is pruned down to the top *W* hits with the highest scores, where *W* is the beam width.

While DNA flows through the pore at an average of 450 bp/s, the electrical signal is recorded at 4000 Hz, leading to roughly 9 measurements/bp on average. Given that we are iterating 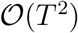 times, doing a paired-read analysis in signal space versus a paired-read analysis in sequence space increases the computational cost ≈ 9 × 9 = 81-fold. To make our calculations more tractable, we make use of an additional heuristic, constraining our search to an alignment envelope[17] of the (*t*_1_, *t*_2_) pairs where the reads are likely to align.

This envelope is estimated by basecalling each read individually, then aligning the two sequences. Reads are basecalled with the Viterbi algorithm, which is faster than beam search with similar performance (Figure 4A), and explicitly returns a signal to sequence mapping, where each nucleotide is mapped to some range of current signal. Then these two basecalled sequences are aligned globally (via Needleman-Wunsch), which generates a mapping of nucleotide from read 1 to nucleotide in read 2, and by extension a range of signal in read 1 to a range of signal in read 2. With some additional padding, this alignment defines a region for our banded 2D beam search (Figure 2).

**Figure 2:**
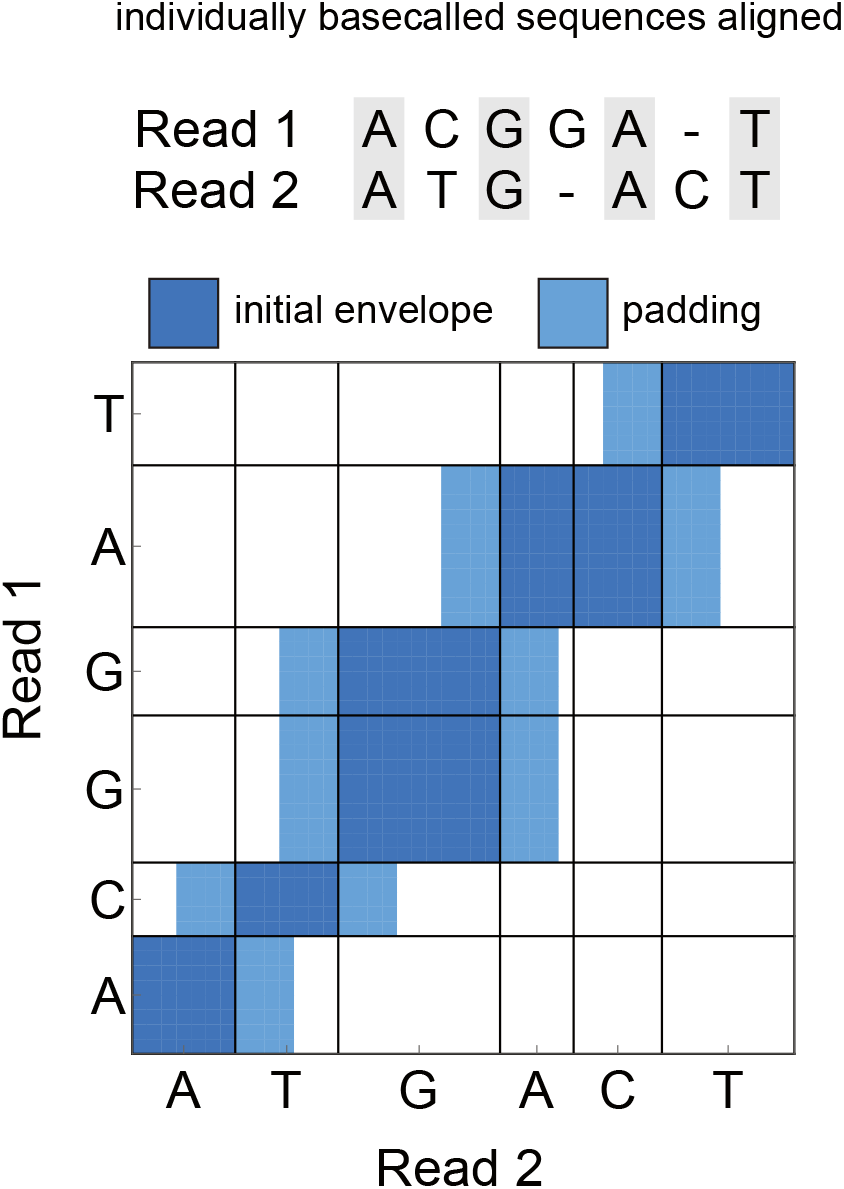
Reads were basecalled individually and the alignment of the resulting sequences was used to construct an alignment envelope in signal space that banded a 2D beam search.

Because the complementary strand passes through the pore with high probability but not every read generates a second strand, there is an additional step during sequencing to determine whether two reads are complementary strands or not. For this work we just relied on the determination of the Guppy basecaller. While one could in theory train two RNNs to operate on the forward and reverse strands and then run consensus on those two probabilities, here we adopt a simpler approach by taking the “reverse complement” of the softmax probabilities, for example:

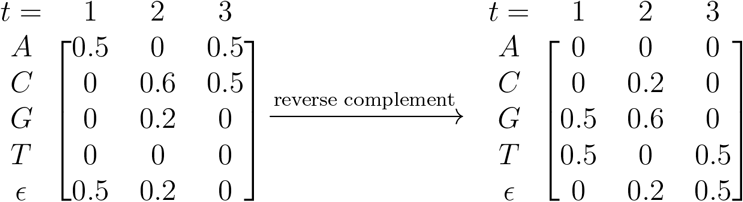

Our test set is a sample of 5,000 R9.4 *E. coli* 1D^2^ read pairs (Oxford Nanopore Technologies, personal communication), comprising 10,000 reads in total. After decoding, reads were aligned to the reference *E. coli* genome with Minimap[19] and the read accuracy is calculated as (number of matches)/(length of alignment).

We test this banded 2D beam search and find that it improves the median accuracy from 86.2% for single reads to 92.1% for read pairs (Figure 3), nearly halving the error rate of our basecaller. While our neural network performs worse than Guppy, which achieved 93.1% 1D accuracy with its larger “high accuracy” network, our consensus algorithm still shows a large median accuracy gain for 1D^2^. Interestingly, the median 1D^2^ accuracy of 94.7% achieved by the “fast” Guppy network exceeded the 94.2% accuracy from the larger “high accuracy” network.

**Figure 3:**
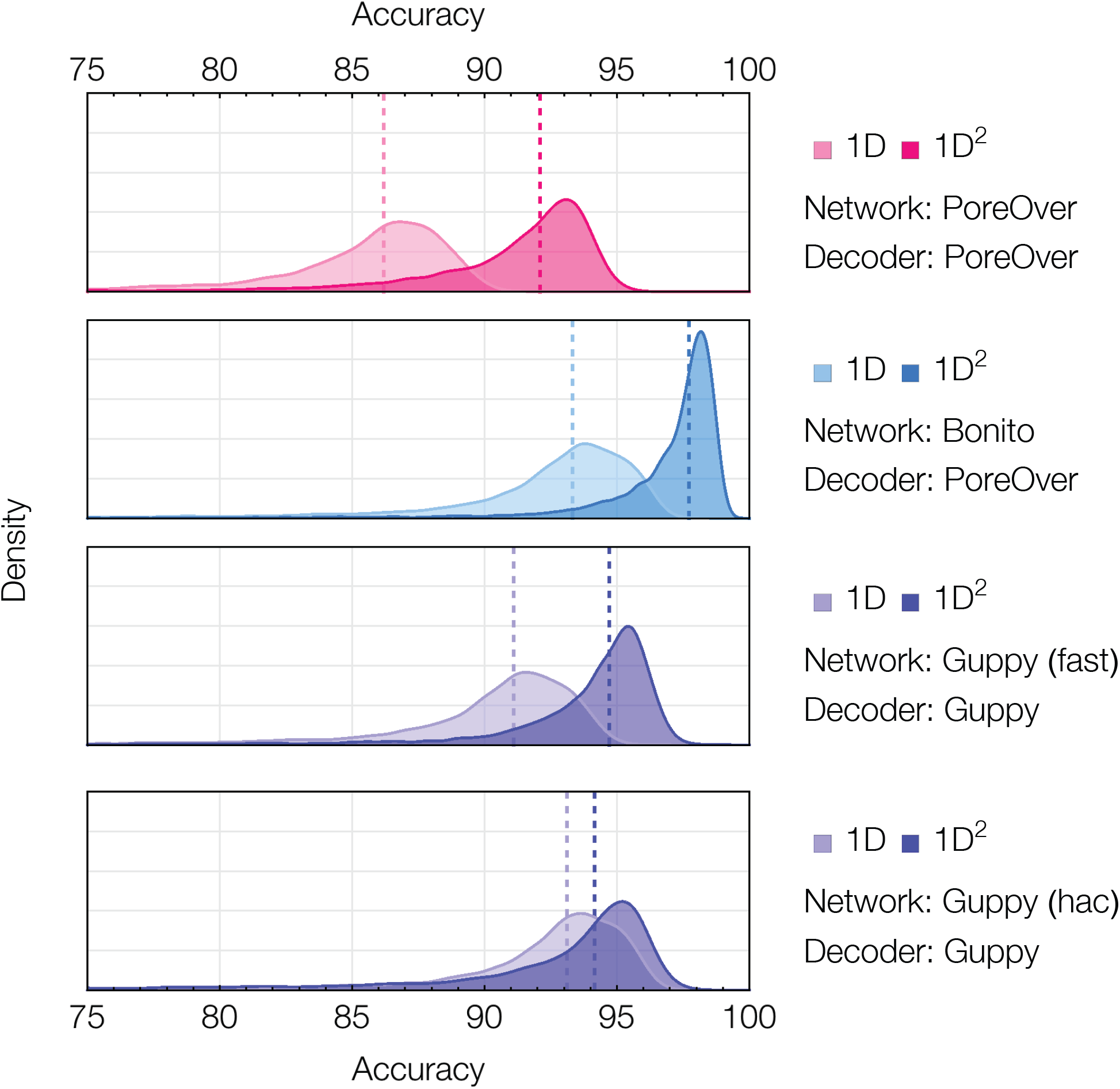
Reads were run through either our PoreOver neural network (magenta) or the Bonito basecaller (blue) to generate softmax probabilities, which were then decoded using our algorithms. Guppy accuracies (in violet) were generated entirely from running the Guppy basecaller and its 1D^2^ basecalling mode without any additional decoding. The Guppy neural network comes with two neural network architectures using either smaller (fast) or larger (high accuracy, hac) recurrent layer sizes. The median accuracy is represented by a dashed line. Bonito version 0.0.1 and Guppy version 3.2.4 were used.

**Figure 4:**
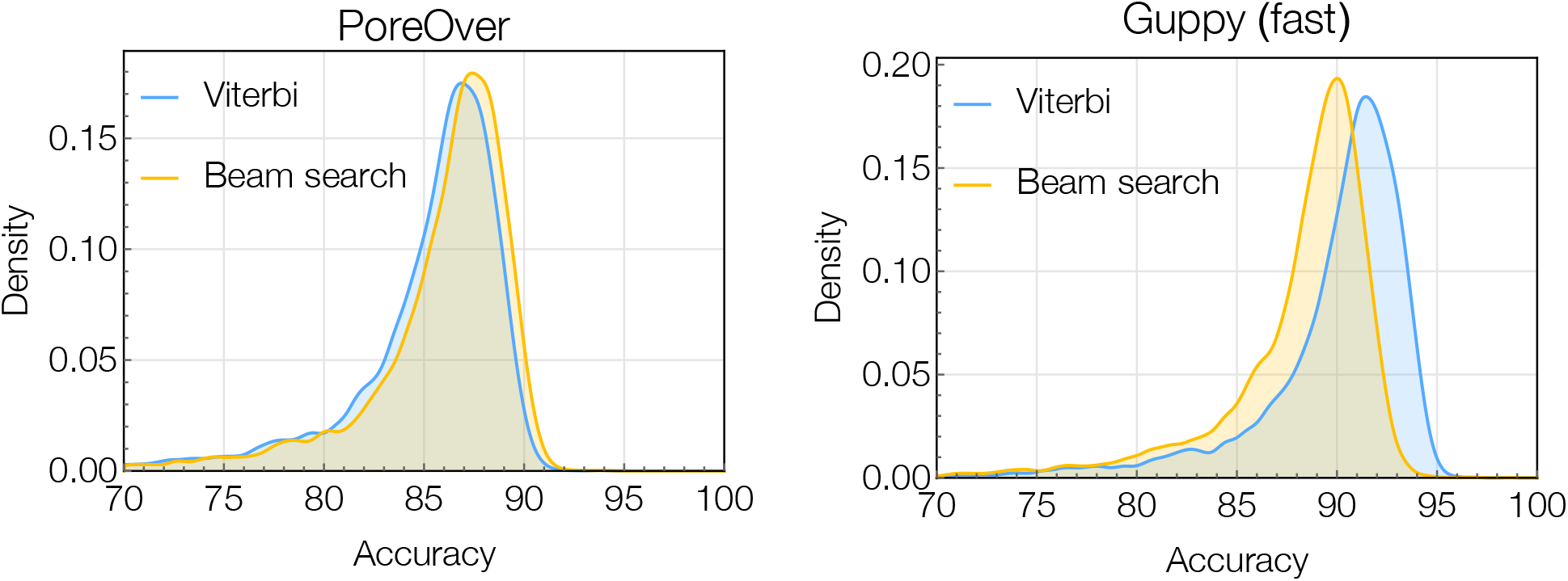
(A) Comparison of single read decoding algorithms using output from our trained network, PoreOver, which implements a simplified CTC model that does not merge repeated characters. (B) Single read decoding algorithms run on output of the Guppy basecaller, which implements a variation of CTC for calling homopolymers called “flip-flop”. In this case the beam search surprisingly returned lower accuracy basecalls than Viterbi decoding. A beam width of 10 was used for both plots.

In December 2019, Oxford Nanopore Technologies released a new research basecaller Bonito, which was inspired by recent successes of purely convolutional neural networks in speech recognition[18]. Bonito is trained using the canonical CTC model (which merges repeated characters), and for each time step generates a probability over each nucleotide plus a gap character. We modified Bonito (version 0.0.1) to save these softmax probabilities and adapted our decoding algorithm to work with this type of CTC model. Using the output of the Bonito basecaller and our consensus algorithm we achieved an accuracy of 97.7%, more than halving the Bonito’s median error rate of 6.7% for single read basecalling and surpassing the consensus accuracy of Guppy’s 1D^2^ method (Figure 3).

### Evaluating decoding algorithms for single read basecalling

We additionally compare single read decoding algorithms using both our own trained basecalling network, as well as ONT’s basecaller Guppy, which implements a variant of CTC called “flip-flop”. The same test set of 10,000 reads was used as in the previous benchmark, though the pairing information was ignored and reads were treated as standard 1D reads.

#### Simplified CTC model

Test reads were split into chunks of 1000 measurements, batched together, and fed through our trained basecalling RNN PoreOver to yield the softmax probabilities. Decoding was then done with both (1) Viterbi best path and (2) a beam search (Figure 4A). The use of beam search over Viterbi yielded a very slight improvement in median accuracy from 86.2% to 86.8%. Interestingly, the beam width had little effect on the overall accuracy, with *W* = 50 yielding nearly identical results. While beam search does yield a slight improvement over Viterbi decoding, it comes at a disproportionately greater computational cost.

#### Flip-flop CTC model

While the earliest nanopore basecallers focused on the task of predicting a sequence of *k*-mers [3], the production basecaller Guppy along with the research basecaller Flappie introduced a character level, CTC-style model known as “flip-flop”. The flip-flop model is an adaptation of CTC for the purpose of better calling homopolymers, a known error mode in nanopore sequencing.

The flip-flop model does not use gaps, as in the standard CTC model but instead has two sets of “flip” (+) and “flop” (−) states, with transitions within flip and flop states only emitting a blank character *ϵ*, *y*(*c, t*) with *c* ∈ {*A*^+^, *C*^+^, *G*^+^, *T* ^+^, *A*^−^, *C*^−^, *G*^−^, *T* ^−^}. Furthermore, transitions from flip to flop states are only allowed between the same nucleotide (e.g. *A*^−^ → *A*^+^). Internally, the basecaller generates a transition matrix for each time step and then runs a Viterbi decoding to generate the final basecalled sequence. The marginalized version of these transition probabilities are stored in FAST5 files, allowing us to use our own decoding algorithms on the flip-flop probabilities. However, we need to adapt the calculation of the forward probabilities to take into account the flip and flop states:

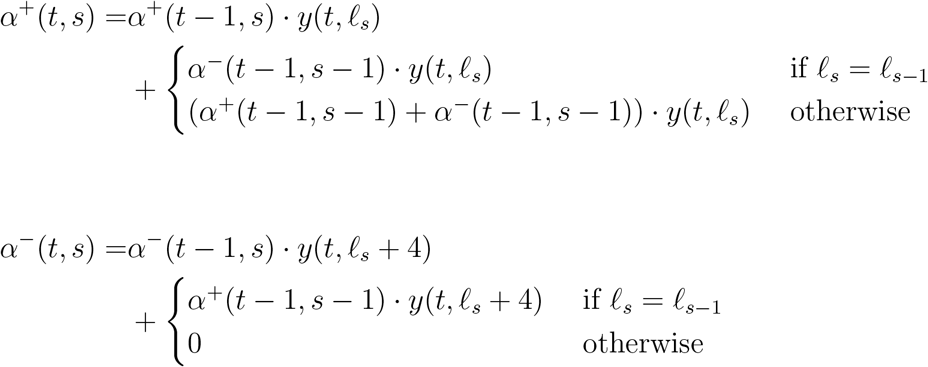

Results from running on the same set of 10,000 reads are shown in Figure 4B. Interestingly for the flip-flop model, the Viterbi algorithm actually outperforms the beam search on this data. The Viterbi decoding yielded a median accuracy of 90.9% while beam search decoding yielded a median accuracy of 88.9%. Despite the sequences returned by the beam search having higher probabilities, they tend to be less accurate when aligned to the reference genome.

## Discussion

In this work, we explored the effects the decoding algorithm has on basecalling accuracy using CTC models. Notably for single read decoding, the Viterbi algorithm is the fastest and performs quite well. Indeed, even for our simplified CTC model, the difference between beam search and Viterbi is quite small and might not be sufficient to justify the increased computational cost.

Curiously with the flip-flop CTC model, the beam search actually did worse than the Viterbi. For some reason, it appears that the more probable sequences aren’t necessarily more accurate. While it is unclear why this was the case with the flip-flop model but not our simplified CTC model, it could be symptomatic of the way these models are trained and evaluated. Much as in several natural language processing tasks, there is a mismatch between the maximum likelihood objective used in training, the CTC loss, and the actual metric used for evaluation, the edit distance or alignment accuracy between the predicted and true labelings. While this metric is discrete and non-differentiable, there has been some success in speech recognition using techniques from reinforcement learning to approximate this gradient and optimize the edit distance directly[12]. Applying policy gradient style approaches that maximize the reward function (in this case alignment accuracy) to CTC basecalling could be an avenue for future research.

In addition to the “flip-flop” model available in the production basecaller Guppy, ONT have also recently introduced another basecalling paradigm known as “run-length encoding, which is implemented in the research basecaller Runnie. Under the run-length encoding model, the neural network outputs the best nucleotide as well as parameters of a discrete Weibull distribution which characterizes the length of the repeat. While this makes single read decoding trivial (by predicting the mode of the parameterized distribution), the 2D beam search described could be adapted to work for run length encoded output. Indeed, one of the strengths of beam search is the ease with which it can be adapted (e.g. to use a language model in speech recognition).

For our consensus application we focused on 1D^2^ sequencing, where our 2D banded beam search yields a significant accuracy improvement. While ONT basecaller Guppy has a built-in 1D^2^ basecalling function, the code is not open source, nor is 1D^2^ basecalling included in research basecallers such as Flappie and Bonito. In contrast, our code is open source and modular in design, making it easily modifiable by the community.

Generalizing beyond a pair of reads, consensus approaches are most relevant to the task of genome assembly polishing. There exist several approaches for multi-read consensus and polishing, a Monte Carlo approach with an HMM model of the pore (nanopolish [20]), or sequence-only approaches using neural networks (Helen+MarginPolish [21] and Medaka^2^). Other work has even used dynamic time warping to generate a consensus at the raw signal level [22].

To our knowledge none of these methods explicitly use the intermediate basecaller probabilities, instead relying largely on the basecalled sequence. While the dynamic programming approach here could be extended for more reads, the curse of dimensionality 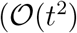 for 2 reads, 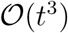 for 3 reads etc.) would necessitate additional heuristics to narrow down the search space. Nevertheless it would be still possible to implement a consensus algorithms (e.g. an MCMC approach) that could exploit the basecaller probabilities for general, multi-read consensus [11].

## Acknowledgements

The authors were supported by NIH/NCI grant CA220441, NIH/NHGRI training grant T32 HG000047, and by Oxford Nanopore Technologies. This research used the computational cluster provided by the Berkeley Research Computing program. We thank Tim Massingham and Marcus Stoiber (Oxford Nanopore Technologies) for helpful discussion.

## Footnotes

1 https://github.com/nanoporetech/tombo

2 https://github.com/nanoporetech/medaka

